# A genome catalog of the early-life human skin microbiome

**DOI:** 10.1101/2023.05.22.541509

**Authors:** Zeyang Shen, Lukian Robert, Milan Stolpman, You Che, Audrey Walsh, Richard Saffery, Katrina J. Allen, Jana Eckert, Angela Young, Clay Deming, Qiong Chen, Sean Conlan, Karen Laky, Jenny Min Li, Lindsay Chatman, Sara Saheb Kashaf, NISC Comparative Sequencing Program, Heidi H. Kong, Pamela A. Frischmeyer-Guerrerio, Kirsten P. Perrett, Julia A. Segre

**Author notes:** These authors contributed equally (Pamela A. Frischmeyer-Guerrerio, Kirsten P. Perrett, Julia A. Segre).

## Abstract

Metagenome-assembled genomes have greatly expanded the reference genomes for skin microbiome. However, the current reference genomes are largely based on samples from adults in North America and lack representation from infants and individuals from other continents. Here we used ultra-deep shotgun metagenomic sequencing to profile the skin microbiota of 215 infants at age 2-3 months and 12 months who were part of the VITALITY trial in Australia as well as 67 maternally-matched samples. Based on the infant samples, we present the Early-Life Skin Genomes (ELSG) catalog, comprising 9,194 bacterial genomes from 1,029 species, 206 fungal genomes from 13 species, and 39 eukaryotic viral sequences. This genome catalog substantially expands the diversity of species previously known to comprise human skin microbiome and improves the classification rate of sequenced data by 25%. The protein catalog derived from these genomes provides insights into the functional elements such as defense mechanisms that distinguish early-life skin microbiome. We also found evidence for vertical transmission at the microbial community, individual skin bacterial species and strain levels between mothers and infants. Overall, the ELSG catalog uncovers the skin microbiome of a previously underrepresented age group and population and provides a comprehensive view of human skin microbiome diversity, function, and transmission in early life.

## Background

In direct contact with the environment, human skin is both a barrier and a habitat for microbes, including bacteria, fungi, and viruses, which help modulate immune responses and provide colonization resistance from adverse species^1, 2^. Skin microbial community composition is shaped both by the ecology of the body site (oily, moist, dry) and skin physiology^1^. For example, during the transition through puberty, the maturation of sebaceous glands creates a lipid-rich environment to facilitate growth of *Cutibacterium*^3^. Compared to adults, early-life skin is characterized by higher water content, lower natural moisturizing factor concentration, and fewer lipids^4, 5^, which provides a distinct cutaneous environment for microbes and a unique habitat to study the skin microbiome.

Human skin microbiota is initially seeded at birth largely from maternal microbiome in association with the mode of delivery^6–8^. This relationship fades within 4-6 weeks^6, 7^, but skin microbial communities at the species level were found to be similar between babies and mothers over weeks to years after delivery^6, 9, 10^. Even though multiple studies have investigated vertical transmission and development of the human gut microbiome^11–14^, mother-to-infant transmission of skin microbiome remains underexplored. Specifically, resolution of vertical transmission of strains on the skin has never been directly demonstrated.

One major challenge in studying the early-life skin microbiome is the lack of microbial reference genomes. Previous skin metagenomic studies found approximately 50% of the metagenomic reads do not match genomes in public databases^1, 15^. Recent advancement in metagenome-assembled genomes (MAGs) has made it possible to generate large genome collections beyond culture-dependent methods^16^. We have recently published the Skin Microbial Genome Collection (SMGC)^17^, which greatly expanded the reference genomes for skin microbiome in adults and substantially improved the classification rate of metagenomic reads. Comprehensive genome collections are also available for human gut microbiome^18–21^. In particular, the recent Early-Life Gut Genomes (ELGG) catalog has indicated great diversity and novelty of early-life gut microbiome compared to later in life^19^. To date, there have been no reports of skin microbial genomes in the first year of life. Comparative research investigating the gut microbiome in different populations also demonstrated great diversity of microbiome in people living in different geographic locations^18, 20, 21^. However, the current skin microbial genomes are derived from mostly adults residing in North America^17^ and lack representation of individuals from other continents.

Here, we sequenced and assembled metagenomes from over 500 skin swabs collected longitudinally at age 2-3 months and 12 months from two body sites of 215 infants born in Australia, providing a catalog of 9,439 nonredundant genomes across multiple kingdoms for early-life skin microbiome. Using these data, we characterized the taxonomic and functional profile of the early-life skin microbiome and investigated the vertical transmission of the skin microbiome between mothers and infants.

## Results

### Deep sequencing of early-life skin metagenomes resulted in 9,439 nonredundant microbial genomes

To obtain comprehensive skin microbiome on early life, we conducted ultra-deep shotgun metagenomic sequencing on 565 skin swabs collected from the cheek and antecubital fossa (inside bend of the elbow) of 215 infants who were part of the VITALITY trial^22^ (Fig. 1a, Supplementary table 1, 2). Among these infants, 69 were sampled longitudinally at 2-3 months and 12 months, 3 were sampled at 2-3 months only, and the rest were sampled at 12 months only. The two skin sites were selected as being representative of sebaceous and moist sites, which are usually inhabited by distinct microbiomes^1^ and have clinical importance for future eczema studies as these are commonly affected sites^23^. Each sample yielded a median of 28.6 million non-human reads (IQR = 11.7-48.6 million). We applied a previously established bioinformatic pipeline^24^ to build MAGs from single samples. To increase MAG quality and the detection of rare species^17^, we pooled reads from the two skin sites within the same individual at each time point to generate MAGs from an additional 283 co-assemblies (Fig. 1a). To generate MAGs, single and pooled sample were assembled with MEGAHIT^25^ and binned with a combination of MetaBAT 2^26^, MaxBin 2^27^, and CONCOCT^28^. Prokaryotic MAGs were refined with metaWRAP^29^ and checked for chimerism with GUNC^30^, while eukaryotic MAGs were checked for quality with EukCC^31^. Eukaryotic viral sequences were detected by aligning the contigs from MEGAHIT to the nucleotide collection database (nt) with BLASTn^32^ and checked for quality with CheckV^33^. After removing redundant genomes across the entire dataset, our analyses yielded 9,194 nonredundant prokaryotic MAGs, 206 nonredundant eukaryotic MAGs and 39 eukaryotic viral sequences, comprising the Early-life Skin Genome (ELSG) catalog.

**Figure 1.**
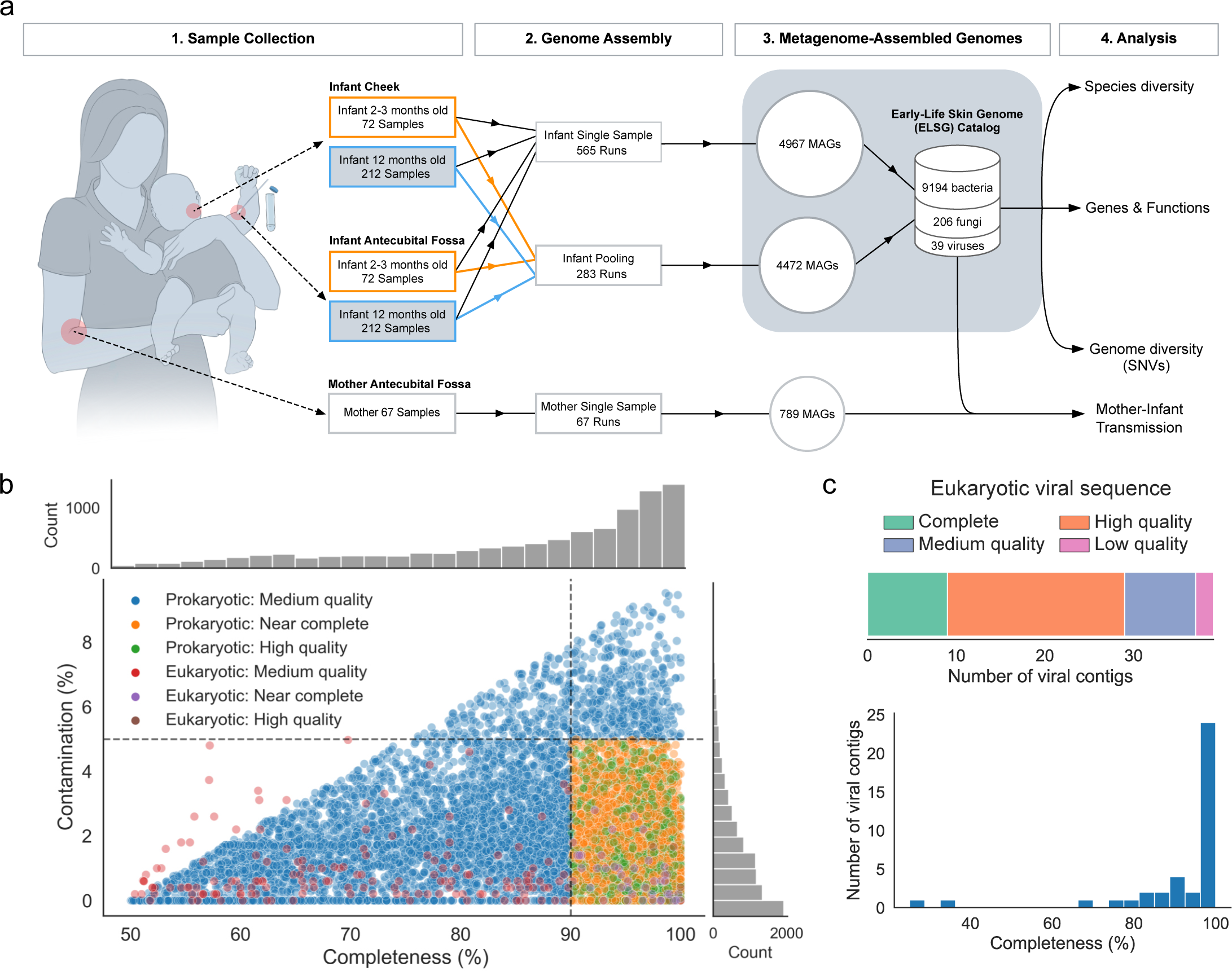
The genome catalog assembled from the early-life skin samples. a. Schematic of study design from sampling to analysis. MAGs were constructed from single samples and pooled samples based on the two body sites of the same infant at each time point. MAGs from infant samples comprise the ELSG catalog. MAGs from mother samples were used for comparative analysis. b. Completeness and contamination for each of nonredundant prokaryotic and eukaryotic MAGs included in the ELSG catalog, colored by the quality level. c. Quality and completeness distribution for eukaryotic viral sequences included in the ELSG catalog.

Among the 9,194 nonredundant prokaryotic MAGs (Fig. 1b, Fig. S1a, Supplementary table 3), 1,550 were classified as “high-quality” (completeness >90%, contamination <5%, and the presence of 5S, 16S and 23S rRNA genes and at least 18 of the standard tRNAs); 2,880 as “near-complete” (completeness >90%, contamination <5%, and didn’t meet the rRNA or tRNA requirement of high-quality MAGs); and 4,764 as “medium-quality” (completeness >50%, contamination <10%, and quality score defined as completeness-5×contamination^18^ > 50) based on the Metagenome-Assembled Genome standard^34^. As a complement to the standard quality metrics, we estimated the level of strain heterogeneity of each MAG using CMSeq^16^ and obtained the median at 0.16% for prokaryotic MAGs. We applied similar criteria to 206 nonredundant eukaryotic MAGs, resulting in 5 “high-quality” (completeness >90%, contamination <5%, and the presence of 5S, 18S, 26S rRNA genes as well as at least 18 of the standard tRNAs), 42 “near-complete” (completeness >90%, contamination <5%, and didn’t meet the rRNA or tRNA requirement of high-quality MAGs), and 159 “medium-quality” MAGs (completeness >50%, contamination <10%) (Fig. 1b, Fig. S1a, Supplementary table 4). Higher quality of MAGs was usually associated with a lower number of contigs, a larger N50, a lower level of strain heterogeneity, a higher read depth, and the presence of more unique tRNAs (Fig. S1a). Among the 39 eukaryotic viral sequences in the ELSG catalog, 9 were classified as “complete” (completeness=100%), 20 as “high-quality” (completeness >90%), 8 as “medium-quality” (completeness >50%), and only 2 as “low-quality” (completeness <50%) according to CheckV^33^ (Fig. 1c, Fig. S1b, Supplementary table 5). Considering the challenge of assembling complete viral sequences from short-read metagenomes^33^, we decided to include the two low-quality sequences in the ELSG catalog.

To investigate transmission, we collected 67 skin swabs from the antecubital fossa of mothers during the 12-month infant visit (Fig. 1a). These samples underwent DNA sequencing and were assembled into individual sample-level MAGs using the aforementioned bioinformatic pipeline. The mother samples yielded a total of 721 bacterial MAGs, 55 fungal MAGs, and 3 eukaryotic viral sequences of medium quality or higher.

### Species diversity in the ELSG catalog

To characterize the phylogenetic diversity of the ELSG catalog, we used 95% average nucleotide identity (ANI) threshold to further cluster the nonredundant MAGs into 1,029 prokaryotic and 13 eukaryotic species-level clusters^35^. We assigned species-level taxonomy to representative prokaryotic MAGs with GDTB-Tk^36^ and to eukaryotic MAGs with >95% ANI to GenBank fungal genomes. Rarefaction analysis showed that the number of species in the ELSG was not saturated, when including MAGs recovered from a single sample. Excluding species recovered from only one sample, which may be transient in nature or individual-specific, the number of species came close to saturation, indicating that the ELSG catalog captured most of the common species present on the early-life skin (Fig. 2a).

**Figure 2.**
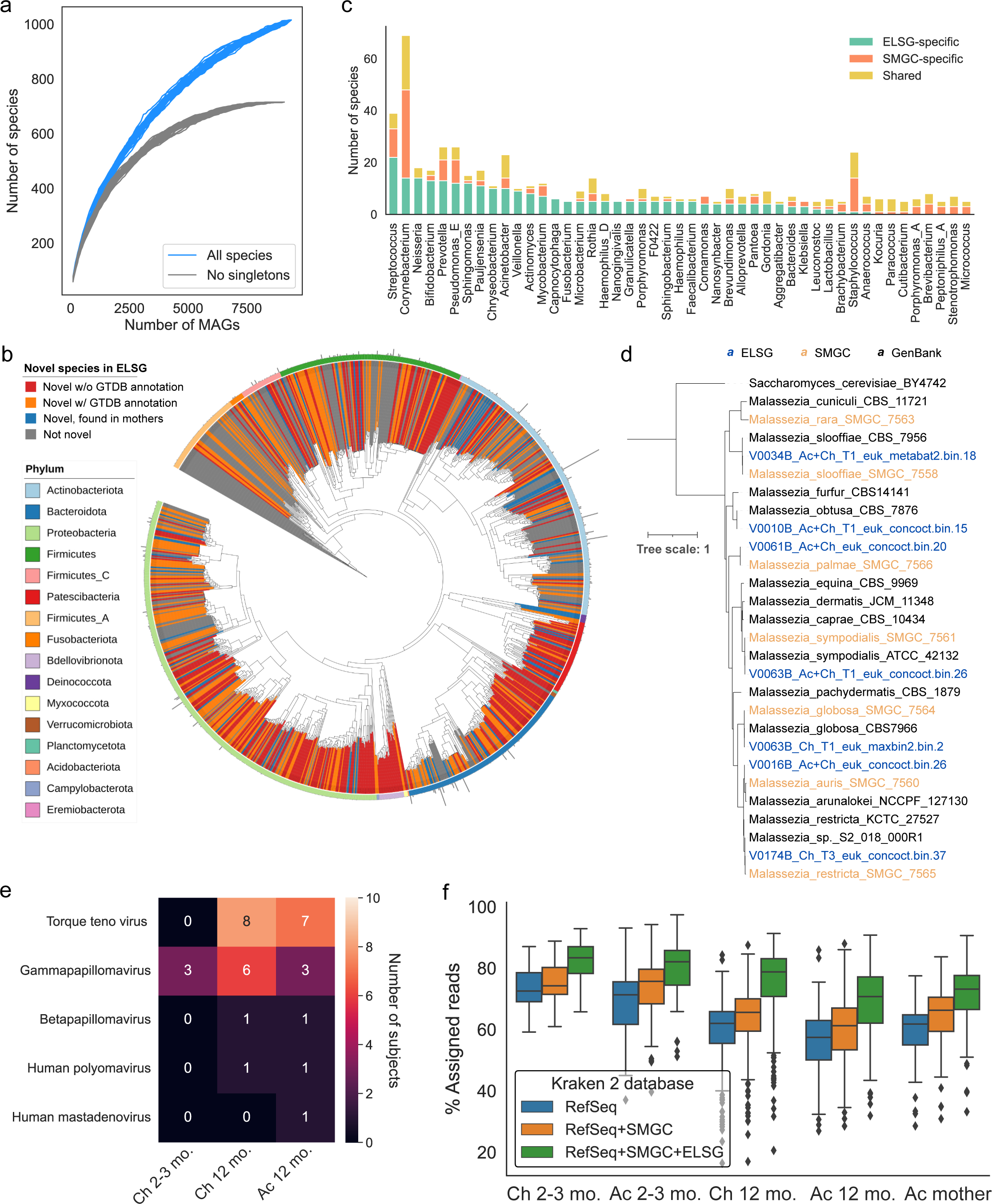
Expansion of species diversity in skin microbiome. a. Rarefaction analysis of the number of species as a function of the number of nonredundant genomes. Curves are depicted both for all the ELSG species and after excluding singleton species (represented by only one genome). b. Phylogenetic tree of the 1,029 representative bacterial MAGs in the ELSG catalog. Clades are colored by GTDB phylum annotation (outer ring) and whether these are novel species (inner shades). Bar graphs in the outermost layer indicate the number of nonredundant genomes within each species-level cluster. c. Comparison of species diversity between the ELSG catalog and the SMGC. Species-level clusters were binned into the genus level in the bar graphs, ordered by a decreasing number of ELSG-specific species. d. Phylogenetic tree of the *Malassezia* genomes from the ELSG and the SMGC together with GenBank reference genomes with *Saccharomyces cerevisiae* as the outgroup. e. Number of infant samples harboring eukaryotic viruses included in the ELSG catalog. f. Proportion of metagenomic reads from skin samples classified with Kraken 2 databases based upon RefSeq, augmented by the SMGC and the ELSG. The boxes represent the interquartile range, and the whiskers indicate the lowest and highest values within 1.5 times the interquartile range.

Next, we explored the novelty of the species diversity in the ELSG catalog. We compared the ELSG catalog with the Skin Microbial Genome Collection (SMGC)^17^, a collection of cultured and uncultured skin microbial genomes primarily based on adult samples in North America, and the Early-Life Gut Genome (ELGG) catalog^19^. Among the 1,029 representative prokaryotic MAGs in the ELSG catalog, 699 clustered independently of any genome from the SMGC and the ELGG, expanding the phylogenetic diversity by 56% (Fig. 2b, Fig. S2a, b, Supplementary table 3). Among these, 313 were not assigned with species-level taxonomy based on GTDB (red in Fig. 2b, Fig. S2b). Note that 79 (11%) species-level clusters overlapped with MAGs built from mothers skin samples (blue in Fig. 2b, Fig. S2b), suggesting these species are likely population-specific rather than early-life-specific. ELSG-specific species spanned 16 different phyla greatly expanding the current knowledge of skin microbiome. Top genera of the early-life-specific species were *Streptococcus*, *Corynebacterium*, *Neisseria*, *Bifidobacterium*, and *Prevotella* (Fig. 2c). Early-life species-level clusters that were also present in the SMGC-specific species were from the genera *Streptococcus*, *Corynebacterium*, and *Prevotella*. As some of the best studied skin genera, *Staphylococcus* harbored very few ELSG-specific species, and similarly *Cutibacterium* species were almost always found in both the ELSG and the SMGC.

Among the eukaryotic genera covered by the ELSG catalog, *Malassezia* was unsurprisingly the dominant genus, followed by *Saccharomyces* (Supplementary table 4), which was not present in the SMGC or mother samples. To compare the *Malassezia* species in the ELSG and the SMGC, we clustered 7 species-level representative MAGs from the ELSG classified to be *Malassezia*, 7 *Malassezia* MAGs from the SMGC, and representative GenBank reference genomes (Fig. 2d). *Malassezia obtusa* was assembled from the early-life skin but not found in the SMGC, whereas *M. rara*, which is a novel species found on the human skin in the SMGC, were not found in the ELSG catalog. Together these findings suggest fungal specificity of early-life skin.

Next, we explored the species diversity of 39 eukaryotic viral sequences in the ELSG catalog. The most prevalent viruses found on infant skin were torque teno virus and gammapapillomavirus (Fig. 2e, Supplementary table 5). Interestingly, the majority of these viral sequences were found exclusively in 12-month infants, except for the gammapapillomavirus discovered on the cheeks of three infants at 2-3 months.

Considering the novel species discovered on early-life skin, we used the ELSG catalog as an additional source of reference genomes to classify shotgun metagenomic reads. By adding the ELSG to a Kraken 2 database^37^ created from the default RefSeq genomes and the SMGC, we obtained a median classification rate of 77% (IQR = 69%-82%) for the early-life skin metagenomic datasets, which was a median of 25% improvement over the standard RefSeq database (Fig. 2f, Fig. S2c). Interestingly, the ELSG also substantially improved the classification rate for metagenomic data of mothers (Fig. 2f, Fig. S2c) and slightly improved read mapping for the antecubital metagenomes of the SMGC (Fig. S2d), suggesting the value of the ELSG in capturing age- or population-specific species.

### Comparison of taxonomic profiles between early-life and adult skin microbiome

We next explored similarities of the infant skin microbial community at two time points as well as the relatedness of infant skin to mothers. The microbial community of infants demonstrated strong skin-site differentiation with cheek and antecubital samples separated on a principal coordinate analysis as well as age differentiation with 2-3 months and 12 months separated for each skin site (Fig. 3a). Interestingly, the microbial community on the antecubital fossa of mothers was most similar to the antecubital fossa of infants at 12 months (Fig. 3a), suggesting a potential trajectory of maturation in the microbial community from early life to adulthood. We calculated Bray-Curtis dissimilarity between the antecubital fossa of babies and mothers and saw a significantly lower beta diversity (p < 1e-4) between related infant-mother pairs compared to unrelated infant-mother pairs, consistent for both infant sexes (Fig. 3b). We also calculated the beta diversity between the two time points of the same infant as compared to different individuals. For both body sites, we saw a significantly lower beta diversity (p < 0.01) within the same individuals, indicating an individualized trajectory of maturation that starts as early as 2-3 months (Fig. S3a). Together, this suggests that the microbial communities on infant skin may be influenced by individual factors, including the mother’s skin microbiome.

**Figure 3.**
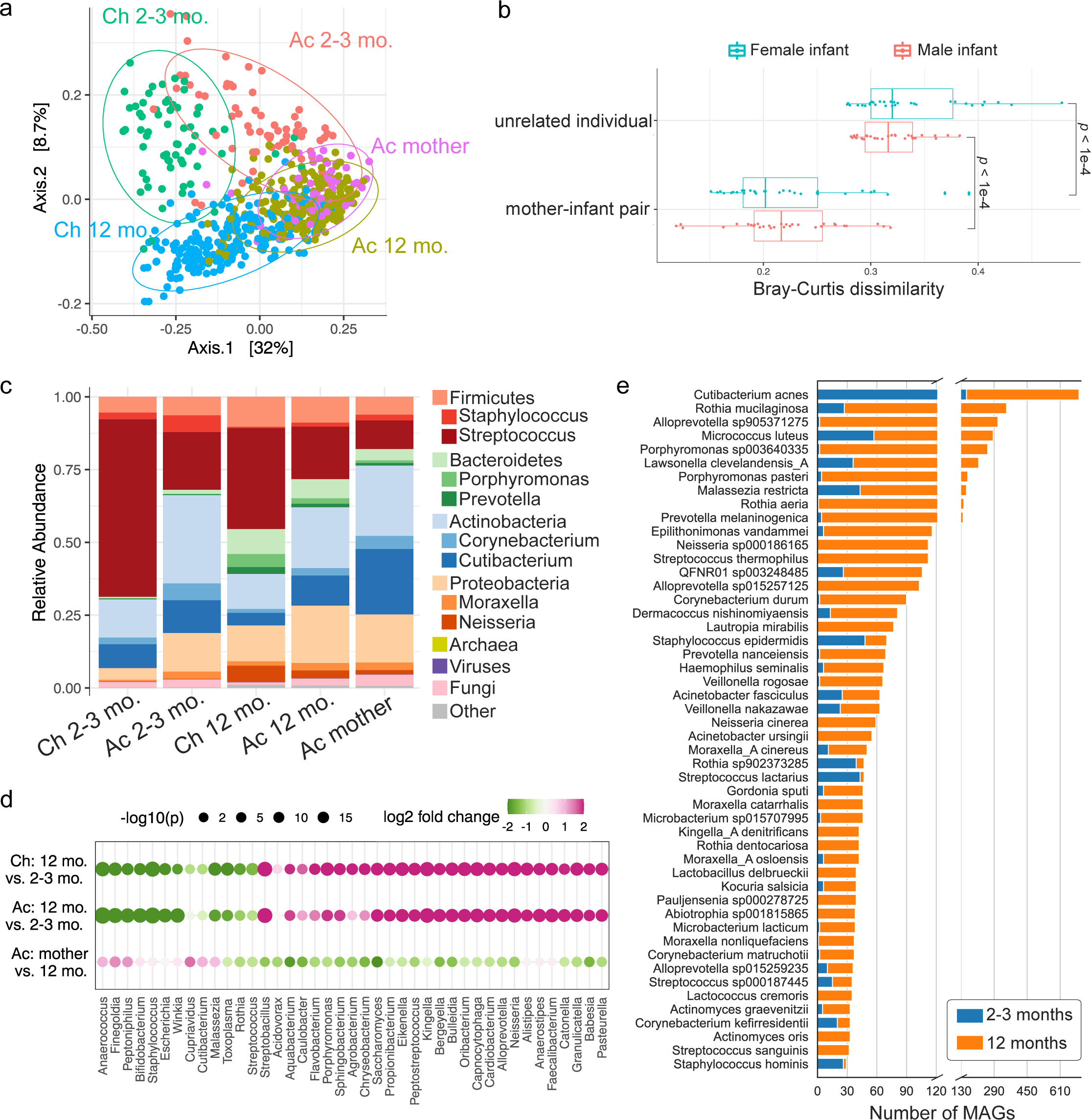
Early-life skin microbial community structure. a. Principal coordinate analysis (PCoA) on Bray-Curtis dissimilarity between the microbial profiles. Each point represents a single sample and is colored by body site and age group. Ellipses represent a 95% confidence interval around the centroid of each sample group. b. The Bray-Curtis dissimilarity of mother-infant pairs comparing related versus unrelated dyads. Median value of each infant and all unrelated mothers was used. Statistical difference was tested by two-sided Wilcoxon rank sum test. c. Relative abundance of skin microbiome averaged for each sample group. Two of the most abundant genera within each bacterial phylum were shown. d. Differential taxa at the genus level between infants of different ages and between infants at 12 months and mothers. The size of the dots represents the log-transformed adjusted p-value from DESeq2, and the color indicates fold changes. The top differentially abundant genera for each comparison were shown. e. Number of species-level MAGs recovered from infants at 2-3 months and 12 months, sorted by the total number of MAGs.

Overall, the skin microbiome of early life contained roughly 97.8% bacteria, 2% fungi, and 0.2% viruses (Fig. 3c), or 92% bacteria, 1% fungi, and 7% viruses after genome size normalization (Fig. S3b). Antecubital fossa of infants generally had a more diverse microbial community than the cheek (Fig. S3c). We also saw an increase in diversity from 2-3 months to 12 months at both body sites (Fig. S3c). At the phylum level, Actinobacteria were more abundant on antecubital fossa, whereas more Firmicutes, particularly *Streptococcus*, was found on cheek (Fig. 3c). Both Bacteroidetes and Proteobacteria gained abundances over time (Fig. 3c). Differential abundance analysis indicated 222 genera significantly (adjusted p < 0.01) gained abundance at antecubital fossa over time and 257 genera increased on the cheek, including *Neisseria* and *Saccharomyces* (Fig. 3d). Another 62 genera and 43 genera lost abundance at 12 months on antecubital fossa and cheek, respectively, including *Staphylococcus* (Fig. 3d), which is consistent with previous studies that also found a decrease in *Staphylococcus* over time^7, 38^. The prevalence of abundant species was correlated with the number of genomes in the ELSG (Fig. S3d). For instance, *Cutibacterium acnes* was the most prevalent species found on early-life skin and contributed the largest number of MAGs in the ELSG (Fig. 3e). Consistent with a higher abundance of *Staphylococcus* at 2-3 months, most of the *Staphylococcus* genomes were assembled from infants at 2-3 months even though the sample size at 2-3 months is much smaller than 12 months (Fig. 3e).

### Comparison of the early-life and adult skin microbiome protein catalogs

To estimate the functional capacity in the ELSG catalog, we predicted protein-coding sequences for each of the 9,194 nonredundant bacterial MAGs, resulting in a total of ∼3.5 million protein clusters at 90% amino acid identity. According to the rarefaction analysis, the protein clusters found in the ELSG catalog were not saturated, but close to saturation when only considering ∼2 million protein clusters that were identified in at least two MAGs (Fig. S4a), consistent with previous findings in gut microbiome^18, 19^. When examining individual species, we discovered that some of the most prominently represented species had either reached a saturation point or were nearing saturation (Fig. 4a). The conspecific gene frequency had a bimodal distribution (Fig. S4b), consistent with observations in the SMGC^17^. We defined those genes shared by at least 90% conspecific genomes of each species as core genes and the rest as accessory genes^18^ (Fig. S4c) and then compared the functions encoded in the core and accessory genes based on several annotation databases. Core genes were generally better annotated than accessory genes in all databases (Fig. S4d). According to COG annotations^39^, core genes were enriched for functions related to metabolism and translation, whereas accessory genes were enriched for functions related to replication, defense mechanisms, and transcription (Fig. 4b). A similar pattern of functional roles performed by core and accessory genes has previously been reported for gut microbiomes^18^.

**Figure 4.**
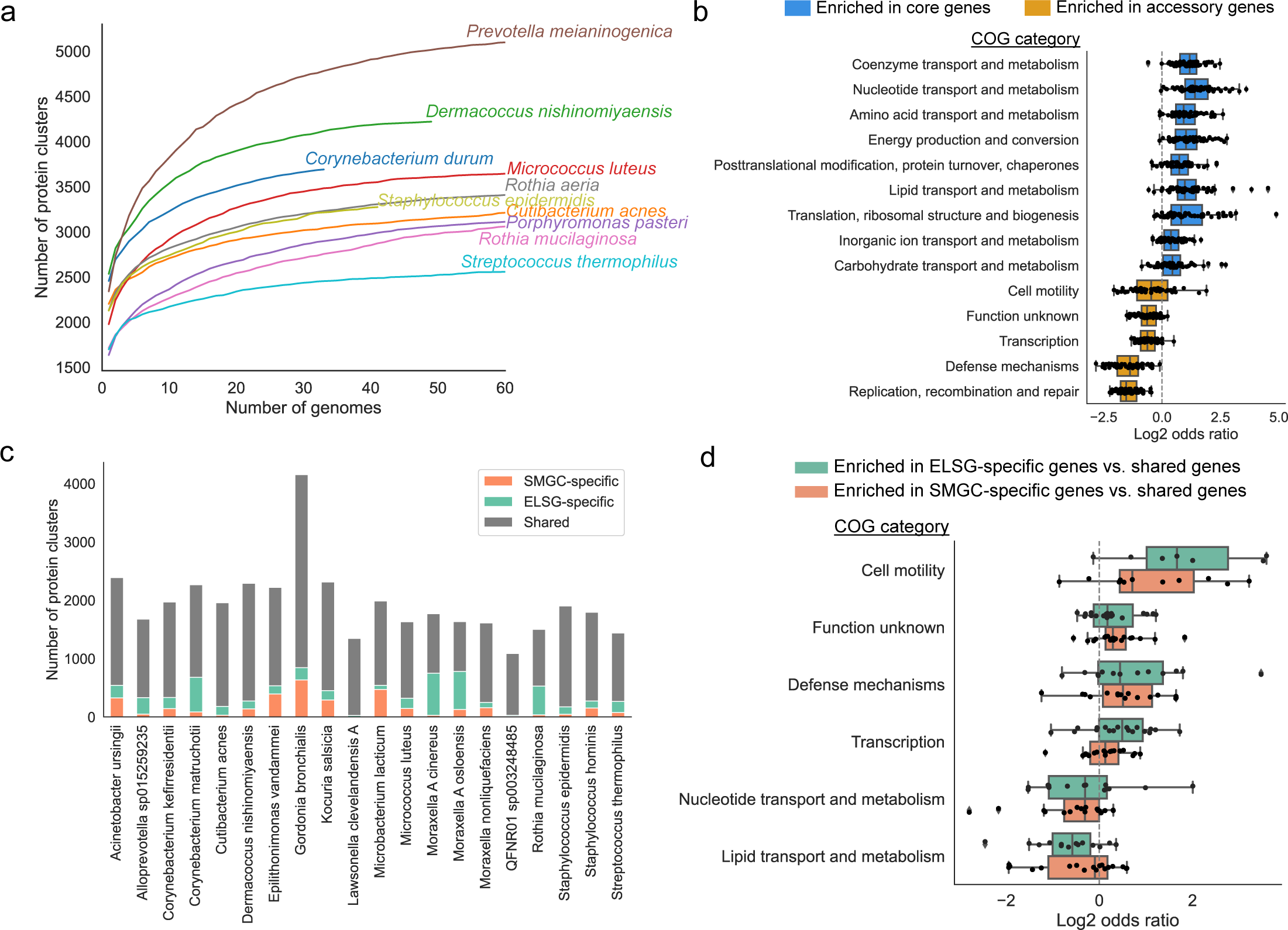
Proteins and functions of early-life skin microbiome. a. Rarefaction curves of the number of protein clusters obtained as a function of the number of species-level genomes. Each curve represents one species. The curves for species with more than 60 genomes are truncated for visualization purpose. b. Comparison of the functional categories assigned to the core and accessory genes for species with at least 10 near-complete or high-quality genomes (>90% completeness, <5% contamination). Each dot represents one species. Odds ratio was calculated from the contingency table with core and accessory genes on one axis and the tested and the other functional categories on the other axis. Only significantly enriched functional categories are shown. Significance was calculated with a two-tailed t-test on log-transformed odds ratios and further adjusted for multiple comparisons using the Bonferroni correction. c. Comparison of the protein clusters between the ELSG and the SMGC for species with at least 5 near-complete or high-quality genomes in each catalog. d. Functional categories enriched in ELSG-specific and SMGC-specific genes compared to shared genes. Each dot represents a species. Only statistically significant categories are shown.

We next compared the pan-genome of early-life skin microbiome with that of SMGC. The pan-genome size was variable between the two genome collections for several species (Fig. S4e). For example, *Micrococcus luteus* had a 14% larger pan-genome in the ELSG catalog, while, in contrast, *Cutibacterium acnes* had a 5% larger pan-genome in the SMGC. Besides the pan-genome size difference, many genes were specific to one collection (Fig. 4c). Interestingly, ELSG- or SMGC-specific genes were enriched in COG categories such as cell motility and defense mechanisms while collection-shared genes were enriched for functions related to metabolism (Fig. 4d).

### Intraspecies genomic diversity between infants and mothers indicates vertical transmission

To characterize the genomic diversity across species-level clusters within the ELSG catalog, we calculated the rate of intraspecies single-nucleotide variants (SNVs). *Rothia mucilaginosa*, a prevalent species on early-life skin, contained the highest SNV density, 40 SNVs per kb, suggesting a great potential of functional variability (Fig. 5a). By contrast, *Cutibacterium acnes*, which was even more prevalent, had a much lower density of only about 5 SNVs per kb. Similarly, *Staphylococcus epidermidis*, another common species found on skin, had about 5 SNVs per kb.

**Figure 5.**
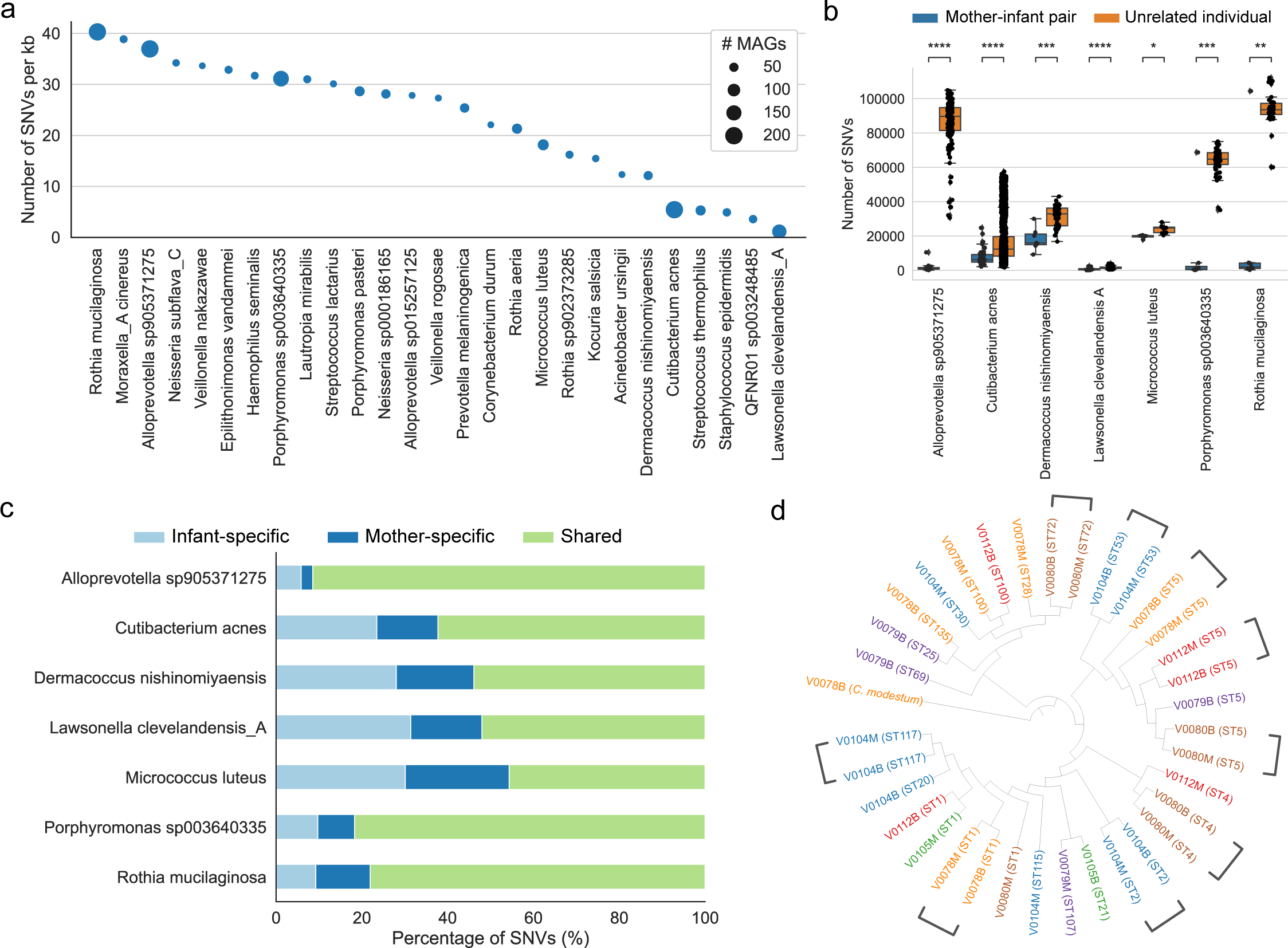
Single-nucleotide variation indicates vertical transmission of skin microbiome. a. Top species with the largest intraspecies SNV density. The size of dots indicates the number of MAGs corresponding to each species. b. Number of SNVs in pairwise comparisons between mother-infant pairs and between infants and unrelated mothers. Only species with genomes from at least 4 mother-infant pairs were considered for analysis. Statistical significance was tested by two-tailed Wilcoxon rank sum test. **P<0.01, ***P<0.001, ****P<0.0001. c. Proportion of SNVs that were found in genomes from infants only or mothers only or both. SNVs were called based on the species-level representative MAG as the reference genome. d. Phylogenetic tree of representative *C. acnes* cultured isolates with *C. modestum* as the outgroup. Source of individual is indicated in the label name and label color. Sequence type is displayed in parentheses.

Next, we compared paired microbial genomes from infants and mothers. For all seven species for which we had MAGs from at least four related infant-mother pairs, there were significantly fewer SNVs genome-wide (p < 0.01) between related infant-mother pairs as compared to unrelated infants and mothers, consistent with vertical transmission of skin microbes between mother and infant (Fig. 5b). By looking at SNVs at protein-coding regions, four of the seven species including *Cutibacterium acnes* had 62% or less SNVs shared by infants and mothers, whereas the other three species including *Rothia mucilaginosa* had over 78% of SNVs shared by infants and mothers (Fig. 5c). The small proportion of age-group-specific SNVs within these three species was also consistent with the strikingly large differences between related and unrelated babies and mothers (Fig. 5b).

Besides the genome sharing between infants and mothers, we also investigated the genome sharing at different ages of infants. For the five species with at least four infants that yielded longitudinal pairs of MAGs, the number of SNVs was generally lower within individuals than across individuals (Fig. S5), suggesting temporally persistent microbial genomes on the host. Due to a limited number of samples, further research is needed to examine the applicability of such observation to a broad spectrum of species.

To further validate the mother-infant transmission of skin microbiome, we cultured *Cutibacterium acnes* from the nasal swabs collected from six pairs of infants and mothers when infants were 12-month-old. Depending on the variable viability of bacteria, we were able to obtain and sequence 4-12 *C. acnes* independent colonies from each individual (Supplementary table 6). Genomes from the related infants and mothers were often closely placed on a phylogenetic tree (Fig. 5d). Consistent with that, we performed multi-locus sequence typing to these genomes and found that four out of six mother-infant pairs shared at least one sequence type (Fig. S5b), which is statistically significant (p = 0.012) based on a permutation test (Fig. S5c). Together, this indicates the mother-infant transmission of skin microbiome is at the strain level.

## Discussion

We present the first genome collection for early-life skin microbiome and the largest skin microbial genome collection to date containing over a thousand species-level clusters of bacterial and fungal genomes and an additional set of eukaryotic viral sequences. To our knowledge, the ELSG catalog is also the first skin microbial genome collection based on samples from Australia. It is an effective resource of genomes to improve the classification of metagenomic reads for early-life samples and geographically distinct studies. The slightly improved classification of North American samples by including the ELSG catalog could be due to the ultra-deep sequencing and the large sample basis of this study, which recovered ultra-rare and low-abundant species present on human skin across continents. Augmented read mapping would be consistent with species that are more highly abundant in infants and at lower abundance in adults. The ELSG catalog includes hundreds of species previously not characterized for skin, many of which are novel species. Considering that skin is still an understudied organ source of microbiome, this study has demonstrated the importance of profiling different age groups and populations to capture a complete catalog of human skin microbiome. Since the ELSG catalog was based on infant samples at age 12 months or less, this resource will be of particular use in studies of childhood cutaneous disorders, such as atopic dermatitis, which commonly begins in infancy.

Our study on the vertical transmission of the skin microbiome was empowered by a substantial number of paired samples collected from infants and mothers. Evidence of vertical transmission was found at the microbial community, individual species, and strain levels. Specifically, infants and their mothers had closely related microbial profiles, relatively similar conspecific MAGs for 7 species, and shared strains of *Cutibacterium acnes*. Likewise, based on the longitudinal samples of infants at 2-3 months and 12 months, we saw evidence of temporal persistence of the microbiome on infant skin in both microbial profiles and genomes. These findings indicate an important role of mothers in shaping the skin microbiome of early life and suggest microbiome at later time point could be affected by what was preceded. Notably, two out of the six mother-infant pairs where we cultured *C. acnes* isolates shared none of their *C. acnes* strains. It suggests that mothers may not be the only source of skin microbiome for infants, and the skin microbial transmission from other sources such as fathers requires further exploration. Further study is also needed to extend our findings to other species that were not investigated in this study.

Based on the ELSG catalog, we analyzed the largest published protein catalog for skin microbiome to estimate the functional capacity. By looking at the conspecific pan-genomes, we summarized the functional categories that distinguish core and accessory genes, which replicated the findings in gut microbiome. Interestingly, genes found only in one of the two current skin genome collections were consistently represented by functions related to defense mechanism and replication, recombination, and repair. These categories are potentially the drivers of functional specificity in early-life skin microbiome. Further experiments are needed to validate the function and importance of individual genes in maintaining homeostasis on early-life skin.

## Conclusions

In summary, our investigation involved profiling the skin metagenomes of infants who had been previously under-represented. This pioneering effort led to the development of the ELSG catalog, which significantly expands the repertoire of skin microbial genomes in infants. The ELSG catalog presents a comprehensive and versatile resource for future studies focused on various aspects of the infant skin microbiome such as microbial transmission and development, and the intricate interplay between disease and the early-life skin microbiota.

## Methods

### Participant recruitment, skin sampling and metagenomic sequencing

New mothers along with their infants were recruited as part of the VITALITY trial^22^. Written informed consent was obtained for all participants in this study. Skin samples were collected from the antecubital fossa and cheek of 72 infants at ages 2-3 months. Sixty-nine of these infants together with 140 additional infants were sampled at the same sites at age 12 months. In addition, 67 of these infants’ mothers were sampled at the antecubital fossa during the same visit when the 12-month samples were taken. To maximize microbial recovery, no bathing was permitted within 24 h of sample collection. Skin was sampled with an established protocol using pre-moistened Puritan foam swabs collected and stored in 100 µL Yeast Cell Lysis Buffer (Lucigen) buffer at -80° and shipped on dry ice. Concomitant with skin sample collection, air swabs were collected as negative controls to account for any potential environmental or reagent contaminants.

Samples were converted to genomic DNA with an established protocol^40, 41^. Briefly, DNA libraries for Illumina sequencing were prepared using the Nextera XT DNA Library Preparation Kit (Illumina) per manufacturer’s instructions with the exception of increasing the AMPure XP Bead clean-up volume from 30 µL to 50 µL. 1 ng of extracted DNA was used as input into the fragmentation step. DNA is simultaneously fragmented and tagged with sequencing adapters in a single-tube enzymatic reaction. Libraries were then sequenced with the Illumina NovaSeq 6000 sequencing platform at the NIH Intramural Sequencing Center for 2 × 150 bp, 50 million paired-end reads per sample.

Most of the negative controls yielded <1% of the reads derived from skin samples except for one. We excluded the skin samples collected at the same time of that air swab together with one infant’s antecubital fossa sample which yielded less than 10,000 reads. Our final set of samples for analysis includes 565 from infants (424 at 12 months (212 infants x 2 skin sites) + 144 at 2-3 months (72 infants x 2 skin sites) - 3 samples failed) and 67 from mothers.

### Bacteria culturing and sequence typing

Nasal culture samples were obtained from infants and mothers during the same visit when infants were 12-month-old using the COPAN eSwab system in 1 mL AMIES and frozen at -80°C. Broths were diluted and plated on Brain Heart Infusion Agar (BHI + 10 µg/mL Fosfomycin) and incubated in an anaerobic chamber for 7 days at 37°C. Colonies were screened with PCR using C. acnes-specific primers PA-1 5’-GGGTTGTAAACCGCTTTCGCTG-3 and PA-2 5’-GGCACACCCATCTCTGAGCAC-3, then streaked for purity on Blood Agar plates (TSA with 5% Sheep Blood – Remel R01201). gDNA was prepared from isolates and sequenced with an established protocol^17^. *C. acnes* genomes were assembled from sequenced reads using SPAdes^42^ and checked for quality using the ‘lineage_wf’ workflow of CheckM v1.1.3^43^. The sequence type of each *C. acnes* genome was identified by multi-locus sequence typing scheme from PubMLST^44^. *C. acnes* genomes of the same individual were first dereplicated at 99.9% ANI with dRep v3.2.2^45^ and then used to build the phylogenetic tree with GToTree v1.6.37^46^ based on the single-copy gene set of Actinobacteria.

### Pre-processing, metagenomic assembly, and contig binning

Metagenomic reads were trimmed for adapters with Cutadapt v3.4 using the parameters “--nextseq-trim 20 -e 0.15 -m 50”^47^ and checked for quality with PRINSEQ-lite v0.20.4 using the parameters “-lc_method entropy -lc_threshold 70 -min_len 50 -min_qual_mean 20 -ns_max_n 5 -min_gc 10 -max_gc 90”^48^. Reads with less than 50 bp length after trimming were removed. The reads were then aligned to the GRCh38 human reference genome with Bowtie2 v2.4.5 using the parameters “--very-sensitive”^49^. The human reads were removed before assembly.

Metagenomic assembly was performed with MEGAHIT v1.2.9 using the default parameters^50^. Pool individual runs were conducted after concatenating the reads from the two skin sites of the same infant at each time point. We performed 283 co-assemblies including 211 from 12 months and 72 from 2-3 months. Contigs were then binned with a combination of MetaBAT 2 v2.15^26^, MaxBin 2 v2.2.7^27^, and CONCOCT v1.1.0^28^ by running the binning module of metaWRAP v1.3.2^29^ with the parameter ‘-l 1500’ indicating the minimal contig length 1500 bp.

### Genome quality assessment

To obtain prokaryotic MAGs, the bins produced by each binning tool were refined with the Bin_refinement module of metaWRAP v1.3.2^29^ using parameters “-c 50 -x 10” enforcing >50% completeness and <10% contamination. The completeness and contamination of refined bins were evaluated with the ‘lineage_wf’ workflow of CheckM v1.1.3^43^. The quality score was calculated as: completeness − 5 × contamination. Ribosomal RNAs in each genome were detected with the ‘cmsearch’ function of INFERNAL v1.1.4 using parameters “--anytrunc -- noali”^51^ against the Rfam covariance models for the 5S (5S_rRNA), 16S (SSU_rRNA _bacteria) and 23S rRNAs (LSU_rRNA _bacteria)^52^. Transfer RNAs of the standard 20 amino acids were identified with tRNAScan-SE v2.0.11 using the parameter ‘-B’ for bacterial species^53^. Each genome was assessed for chimerism with GUNC v1.0.5^54^. The MAGs with contamination greater than 0.05, clade separation greater than 0.45 and a reference representation score greater than 0.5 were excluded. Based on the Metagenome-Assembled Genome standard^34^, MAGs with >90% completeness, <5% contamination, the presence of 5S, 16S and 23S rRNA genes, and at least 18 tRNAs were reported as high-quality draft genomes. MAGs with >90% completeness and <5% contamination but missing the rRNAs or tRNAs were reported as near-complete genomes. MAGs with >50% completeness and <10% contamination were reported as medium quality.

To assess eukaryotic MAGs, the bins from the three binning tools were estimated for completeness and contamination with EukCC v2.1.0^31^. rRNAs and tRNAs were identified using the same approach above except that the Rfam^52^ covariance models 5_S_rRNA, SSU_rRNA_eukarya and LSU_rRNA_eukarya were used to find 5S, 18S and 26S, respectively. Bins with >90% completeness, <5% contamination, the presence of 5S, 18S and 26S rRNA genes, and at least 18 tRNAs were reported as high-quality draft genomes. Those with >90% completeness and <5% contamination but not satisfying the rRNAs and tRNAs requirements were defined as near-complete. The rest of the bins with >50% completeness and <10% contamination were reported as medium-quality genomes.

We further mapped each contig of MAGs to the nt database with BLASTn v2.8.0^32^ to assess viral contamination. Contigs with the top hit of a eukaryotic viral genome with >95% nucleotide identity, >1000 bp aligned sequence, and >70% total contig aligned were removed. The contig number and N50 of MAGs were calculated using in-house scripts. Read depth was calculated by first mapping the raw reads back to MAGs Bowtie2 v2.4.5^49^ using the default parameters and then calculating mean depth with SAMtools v1.16.1^55^. The strain heterogeneity was estimated by the “polymut.py” script of CMSeq v1.0.4 with parameters “--mincov 10 --minqual 30--dominant_frq_thrsh 0.8”^16^.

Eukaryotic viral sequences were detected by aligning the contigs from MEGAHIT to the nt database with BLASTn v2.8.0^32^ by requiring >90% nucleotide identity, >1000 bp aligned sequence, and >70% total contig aligned. The quality of viral sequences was assessed with CheckV v1.0.0 based on database v1.5^33^.

### Redundancy removal and species clustering

To remove redundant genomes that were recovered by both single and pooled sample runs, we dereplicated MAGs at a 99.9% ANI threshold with dRep v3.2.2 using parameters ‘-pa 0.999 -- SkipSecondary -cm larger --S_algorithm fastANI -comp 50 -con 10’^45^. fastANI v1.33^35^ was used to accelerate the process. Dereplication was performed on prokaryotic MAGs and eukaryotic MAGs separately.

The MAGs were clustered at the species level by dereplicating at a 95% ANI threshold with dRep v3.2.2 using parameters ‘-pa 0.90 -sa 0.95 -nc 0.30 -cm larger --S_algorithm fastANI -comp 50 -con 10 --run_tertiary_clustering --clusterAlg single’^45^. Representative genome of each species-level cluster was selected based on the dRep scores derived from genome completeness, contamination, strain heterogeneity, and contig N50.

### Taxonomic assignment and phylogenetic analysis

Taxonomic annotation of prokaryotic MAGs was assigned with the “classify_wf” workflow of GTDB-Tk v2.1.0 using default parameters and GTDB database release 207^36, 56^. The phylogenetic tree of bacterial representative genomes of species-level clusters was built with IQ-TREE v1.6.12 using the parameter “-m MFP”^57^ based on the protein sequence alignments generated by GTDB-Tk.

The eukaryotic MAGs were compared with all of the GenBank fungal genomes first using Mash^58^ and then assigned species-level taxonomy with at least 95% ANI calculated by fastANI v1.33^35^. The phylogenetic tree was built with the script BUSCO_phylogenomics.py (https://github.com/jamiemcg/BUSCO_phylogenomics) based on single-copy marker genes identified by BUSCO v4.1.3 using the parameter “-m geno -f --auto-lineage-euk”^59^. The phylogenetic trees were visualized with iTOL^60^. The taxonomic classifications of viral sequences were assigned by the top alignment hit from BLASTn^32^.

### Metagenomic read classification and microbial abundance estimation

Metagenomic reads were mapped with Kraken v2.1.2 using parameters “--confidence 0.1 -- paired”^37^ against the standard RefSeq database (release 99) and two custom database with additional representative genomes from the SMGC and ELSG catalogs. To integrate the genome catalogs with the RefSeq genomes, we first converted GTDB taxonomy to NCBI taxonomy using the “gtdb_to_ncbi_majority_vote.py” script available in the GTDB-Tk repository^36^ and then obtained NCBI taxonomy IDs corresponding to the species- and genus-level taxonomy of each genome with taxonkit v0.12.0^61^. We excluded 22 and 106 representative MAGs from the SMGC and the ELSG, respectively, which did not have a match ID at the genus level. For MAGs with a match ID at the genus level but not at the species level, we created a new taxonomy ID associated with each MAG when building the Kraken databases. Classification improvement was calculated on a per-sample basis as (proportion of reads classified with custom database − proportion of reads classified with RefSeq database)/proportion of reads classified with RefSeq database × 100. Species-level microbial abundances were computed with Bracken v2.5 using parameters “-r 100 -l S”^62^.

### Alpha and beta diversity calculation

Skin metagenomic data with less than 800,000 classified reads were excluded (4% of samples). The remaining samples were first rarefied and then calculated for the number of species with ≥5 reads (richness) and Shannon index with the “diversity” function of vegan package in R v4.1. To calculate the beta diversity, we first removed taxa present in ≤20% samples and then performed log transformation on species abundances after adding pseudocount 1. Bray-Curtis dissimilarity was calculated with the “distance” function of phyloseq v1.38.0^63^ in R v4.1. Principal coordinate analysis was conducted based on Bray-Curtis dissimilarity with the “ordination” function of phyloseq package.

### Differential abundance analysis

Differential abundance was calculated with DESeq2 v1.34.0^64^ using the parameters ‘test=“Wald”, sfType=“poscounts”, fitType=“local”’ based on the rarefied raw reads as used for diversity calculation. Low-prevalence taxa present in less than 10% of samples were removed. Comparisons were conducted for each of the two skin sites, comparing 2-3 months and 12 months; and for antecubital fossa, comparing infants at 12 months and mothers. Significantly differential taxa were identified by <0.01 adjusted p-value and >2-fold change.

### Pan-genome analysis and functional annotation

Protein-coding sequences (CDS) of each genome were predicted and annotated with Prokka v1.14.6 using parameter “--kingdom Bacteria”^65^. Protein clustering across all species of genomes was conducted with the ‘easy-linclust’ function of MMseqs2 v13-45111 using parameters ‘--cov-mode 1 -c 0.8 --kmer-per-seq 80 --min-seq-id 0.9’ to generate protein clusters at 90% amino acid identity, respectively^66^.

The pan-genome analysis was performed only on near-complete and high-quality genomes. Species with at least ten near-complete or high-quality nonredundant genomes were analyzed with Panaroo v1.3.0 using the parameters ‘--clean-mode strict --merge_paralogs -c 0.90 -- core_threshold 0.90 -f 0.5’ for ≥90% amino acid identity and a family threshold of 50%^67^. Functional annotation of all protein sequences was performed with eggNOG-mapper v2.1.6^68^ to obtain COG^39^, KEGG^69^, Pfam^70^ and GO^71^ annotations.

### SNV analysis

To assess SNV density of species, we first mapped conspecific genomes to the representative genome using the ‘nucmer’ program of MUMmer v3.1^72^, filtered alignments with the ‘delta-filter’ program using parameters ‘-q -r’, and then identified SNVs with the ‘show-snps’ program. SNV density of each genome was computed by dividing the number of SNVs by the size of the representative genome. Only SNVs which occurred in at least two conspecific genomes were included in the analysis. The final SNV density of each species was the mean of SNV densities of all conspecific genomes. The same programs and parameters were used for mother-infant genome comparisons.

### Statistical analysis

Statistical analyses were performed using ggpubr package in R v4.1 or scipy package in Python v3.9.9. Two-sided Wilcoxon rank sum tests and t-tests were used to evaluate differences between groups. Pearson correlation coefficient was used to assess correlation. Functional enrichment analysis was performed using two-sided Fisher’s exact test, with p-values adjusted by the Bonferroni method. The permutation test (n = 1,000) was applied to assess the significance of sequence type sharing between mothers and infants.

## Availability of data and materials

The raw metagenomic sequencing data are available in the NCBI BioProject database under project number PRJNA971252. The MAGs of the ELSG catalog can be found at https://research.nhgri.nih.gov/projects/ELSG/. Additionally, on the same website, users have access to download nonredundant genomes, species-level representative genomes, phylogenetic tree files, protein catalog, pan-genome annotations, and a custom Kraken 2 database based on the ELSG catalog. All the code utilized in this study is available on GitHub at https://github.com/skinmicrobiome/ELSG.

Other publicly available data used in this project: SMGC is available at http://ftp.ebi.ac.uk/pub/databases/metagenomics/genome_sets/skin_microbiome. Shotgun metagenomic sequencing data used in the SMGC is accessed from the NCBI Sequence Read Archive under accession number SRP002480. ELGG catalog is available at https://doi.org/10.5281/zenodo.6969520.

## Acknowledgements

Research reported in this publication was performed in part as a project of the Immune Tolerance Network and supported by the National Institute of Allergy and Infectious Diseases (NIAID) of the National Institutes of Health (NIH) under Award Number UM1AI109565. The study made use of the computational resources provided by the NIH HPC Biowulf Cluster (http://hpc.nih.gov) and received support from the National Health and Medical Research Council (NHMRC) of Australia, the Intramural Research Programs of the National Human Genome Research Institute, the NIAID as well as the National Institute of Arthritis and Musculoskeletal and Skin Diseases. The authors are grateful to the families for participation.

---

**Figure S1.** Quality metrics of the nonredundant genomes in the ELSG catalog.

a. Genomes were stratified by quality level with colors matching those in Figure 1. Box lengths represent the interquartile range, and whiskers indicate the lowest and highest values within 1.5 times the interquartile range.

b. Eukaryotic viral sequences were stratified by quality level with colors matching those in Figure 1.

**Figure S2.** Expansion of species diversity in the ELSG catalog.

a. Comparison of species-level representative MAGs from three genome catalogs: ELSG, SMGC, and ELGG. The numbers indicate MAGs from each catalog that did or did not cluster with other catalogs at the species level.

b. Phylogenetic diversity accounted by known and novel species. Colors match Figure 2.

c. The improvement rate of read classification over the standard Kraken 2 RefSeq database.

d. Kraken 2 classification rate of reads from published skin metagenomic data.

**Figure S3.** Early-life skin microbial community structure.

a. The Bray-Curtis dissimilarity of microbial community between two time points of the same infant compared with that of different infants. Median value of each infant and all other infants was used to plot. Statistical difference was tested by two-sided Wilcoxon rank sum test.

b. Relative abundance of skin microbiome averaged for each sample group after genome size normalization, which emphasized the viral community.

c. Alpha diversity (richness and Shannon index) of skin samples divided by age group and skin site.

d. Relationship between species prevalence and number of MAGs. Each dot represents a species, the color of which indicates maximum relative abundance among all samples. The prevalence of a species was calculated as the number of samples with >0.1% relative abundance of that species. Pearson’s correlation coefficient and p-value indicate a significant correlation.

**Figure S4.** Proteins and functions of early-life skin microbiome.

a. Rarefaction curves of the number of protein clusters as a function of the number of genomes for all species combined. Separate curves are depicted for proteins clustered at 90% amino acid identity for all protein clusters and after excluding singleton protein clusters (containing only one protein sequence).

b. The number of genes in relation to the fraction of conspecific genomes where genes were found. Only near-complete and high-quality genomes were considered in the analysis.

c. Number of genes shared by conspecific genomes in relation to the cutoff on the fraction of conspecific genomes. Vertical dashed line represents the 90% threshold used in this study to define core genes.

d. Proportion of core and accessory genes annotated by various databases including Clusters of Orthologous Genes (COG), Kyoto Encyclopedia of Genes and Genomes (KEGG), Pfam, and Gene Ontology (GO). Each dot represents a species. Only species with at least 10 near-complete or high-quality genomes are included. Statistical significance was tested by two-tailed Wilcoxon rank sum test.

e. Comparison of the rate of gene gain between the ELSG and the SMGC. Only near-complete and high-quality genomes are included. Dashed line connects the end point of the collection with fewer number of genomes of the same species.

**Figure S5.** Intraspecies single-nucleotide variation and vertical transmission.

a. Number of SNVs in pairwise comparisons between genomes of the same infant at 2-3 months and 12 months (same infant) and between any two genomes assembled from different times (different infant). Only species with genomes from at least 3 infants at different times were considered for analysis. Statistical significance was tested by two-tailed Wilcoxon rank sum test. *P<0.05, **P<0.01. Non-significance (“ns”) indicates P>0.05.

b. Number of sequence types of *C. acnes* cultured isolates from each individual.

c. Average number of shared sequence types of *C. acnes* cultured isolates between related infants and mothers (orange dashed line) and between any two individuals after 1,000 permutations (histogram).

