## Supplementary figures and images for "A genome catalog of the early-life human skin microbiome"

### Supplementary Figures

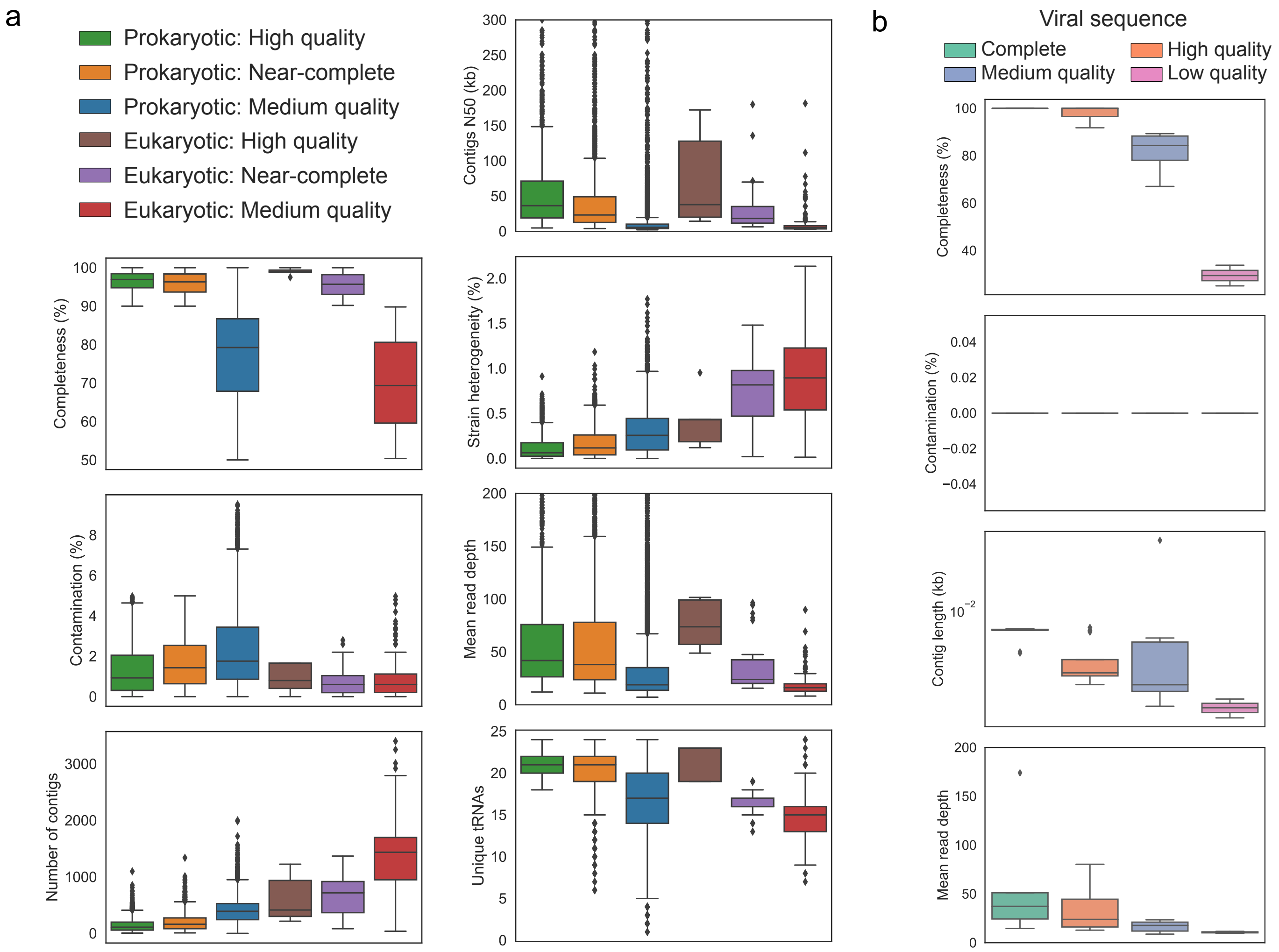

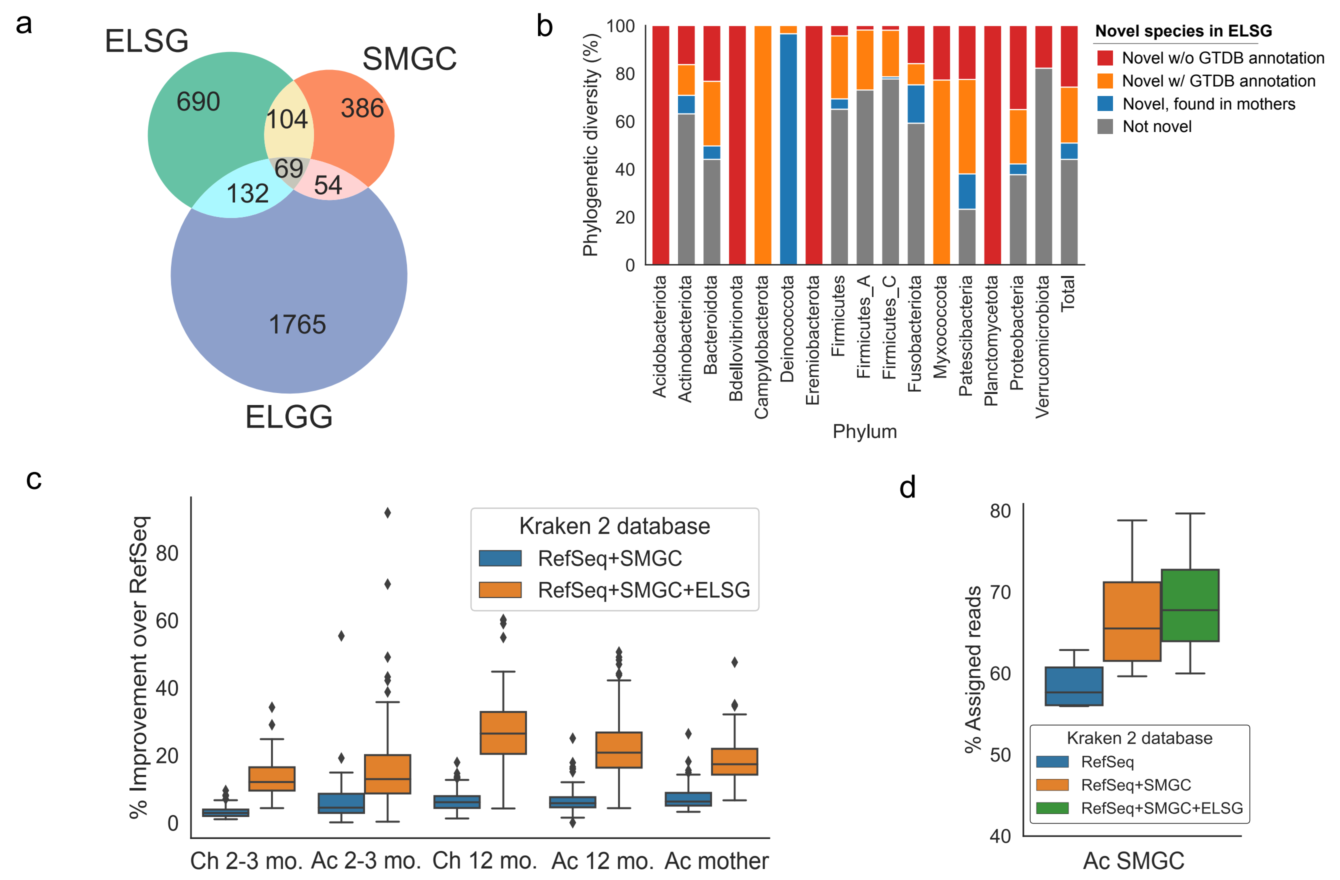

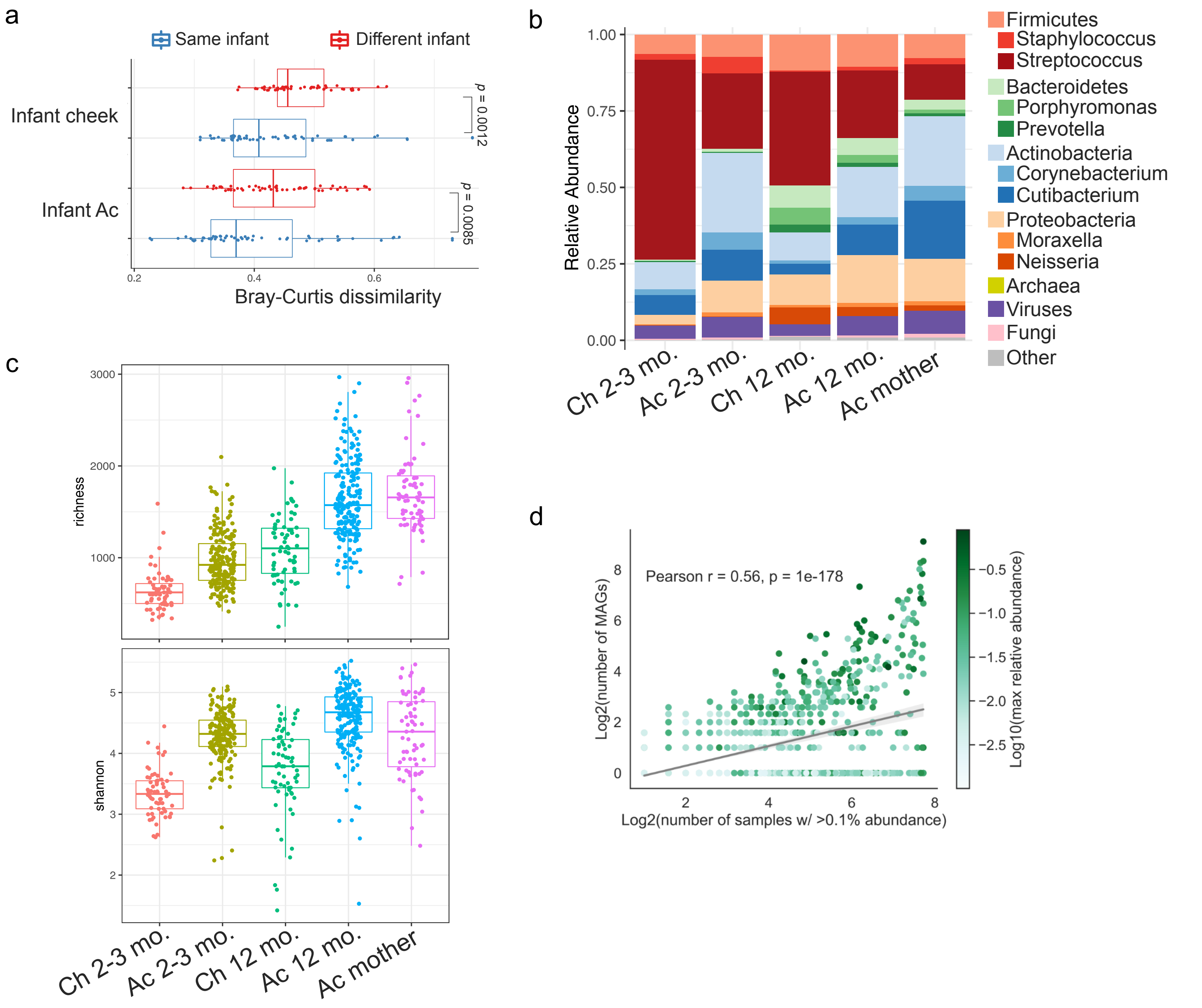

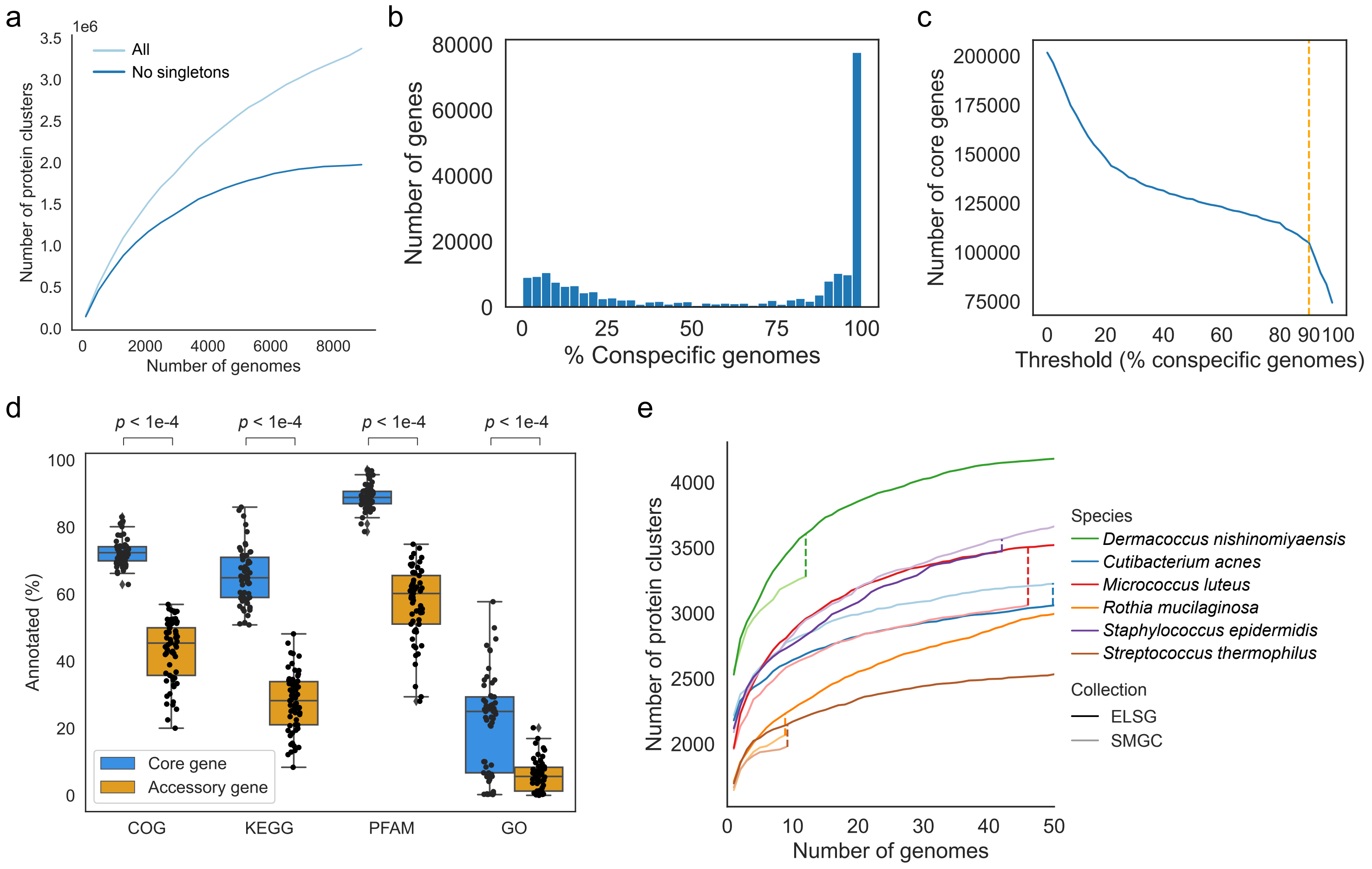

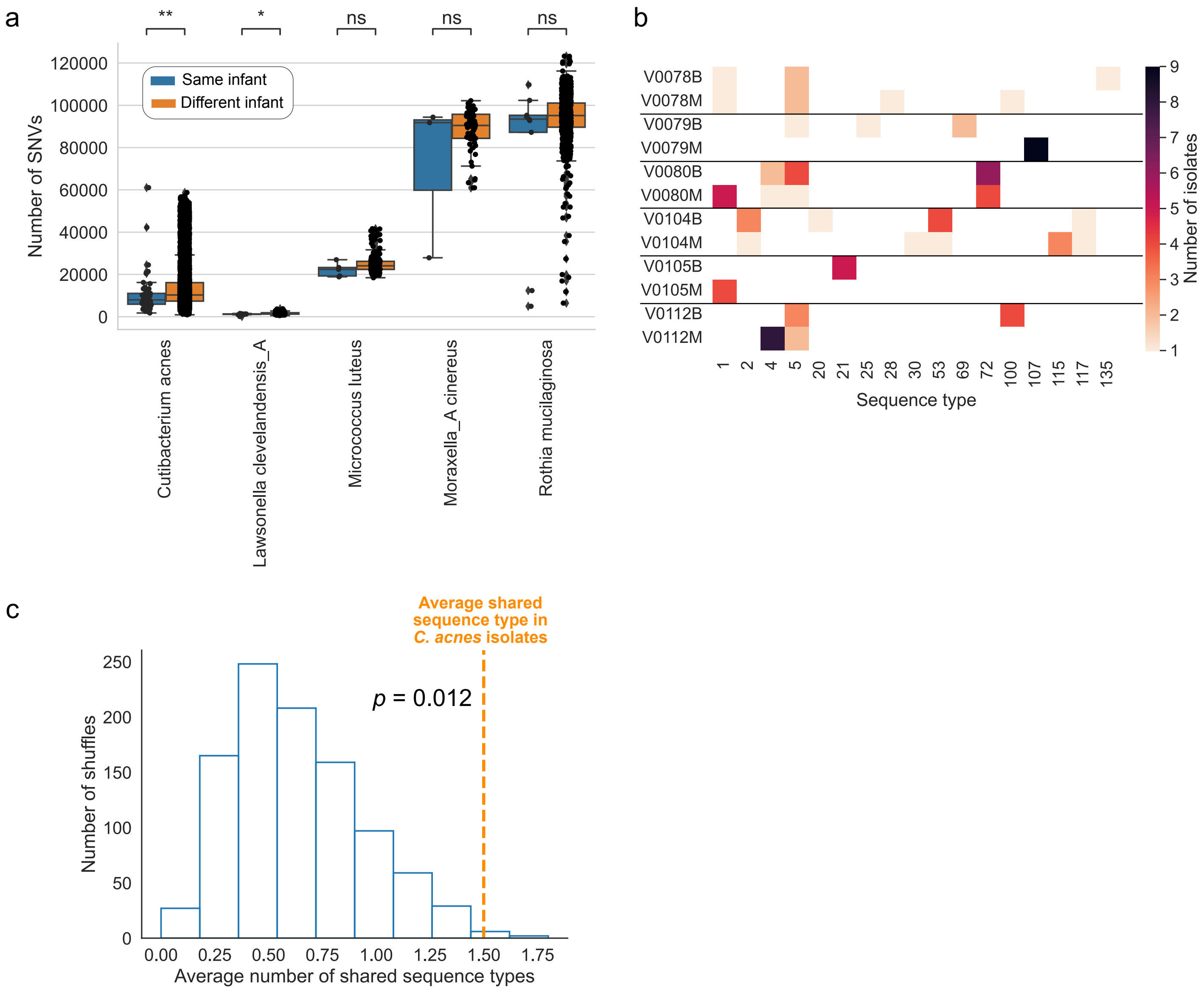
